# Characterising the mechanisms underlying genetic resistance to amoebic gill disease in Atlantic salmon using RNA sequencing

**DOI:** 10.1101/699561

**Authors:** Diego Robledo, Alastair Hamilton, Alejandro P. Gutiérrez, James E. Bron, Ross D. Houston

## Abstract

**Background:** Gill health is one of the main concerns for Atlantic salmon aquaculture, and Amoebic Gill Disease (AGD), attributable to infection by the amoeba *Neoparamoeba perurans*, is a frequent cause of morbidity. In the absence of preventive measures, increasing genetic resistance of salmon to AGD via selective breeding can reduce the incidence of the disease and mitigate gill damage. Understanding the mechanisms leading to AGD resistance and the underlying causative genomic features can aid in this effort, while also providing critical information for the development of other control strategies. AGD resistance is considered to be moderately heritable, and several putative QTL have been identified. The aim of the current study was to improve understanding of the mechanisms underlying AGD resistance, and to identify putative causative genomic factors underlying the QTL. To achieve this, RNA was extracted from the gill and head kidney of AGD resistant and susceptible animals following a challenge with *N*. *perurans*, and sequenced.

**Results:** Comparison between resistant and susceptible animals pointed to differences mainly in the local immune response in the gill, involving red blood cell genes and genes related to immune function and cell adhesion. Differentially expressed immune genes highlighted differences in the Th2 and Th17 responses, which are consistent with the increased heritability observed after successive challenges with the amoeba. Five QTL-region candidate genes showed differential expression, including a gene connected to interferon responses (*GVINP1*), a gene involved in systemic inflammation (*MAP4K4*), and a positive regulator of apoptosis (*TRIM39*). Analyses of allele-specific expression highlighted a gene in the QTL region on chromosome 17, cellular repressor of E1A-stimulated genes 1 (*CREG1*), showing allelic differential expression suggestive of a cis-acting regulatory variant.

**Conclusions:** In summary, this study provides new insights into the mechanisms of resistance to AGD in Atlantic salmon, and highlights candidate genes for further functional studies that can further elucidate the genomic mechanisms leading to resistance and contribute to enhancing salmon health via improved genomic selection.

## BACKGROUND

Gill health is currently one of the major concerns for Atlantic salmon farming worldwide. Fish gills are multifunctional organs fundamental for gas exchange, ionoregulation, osmoregulation, acid-base balance and ammonia excretion, but also play an important role in hormone production and immune defence (Rombough 2007). Gills are constantly exposed to the marine environment, and are often the first line of defence against pathogens. Gill damage is often observed in Atlantic salmon under farming conditions, and can pose a significant welfare, management and economic burden. While the aetiology of gill disorders is complex, Amoebic Gill Disease (AGD) is currently regarded as a key threat to gill health. This disease adversely affects the gill, and can result in respiratory distress, and ultimately mortalities if left untreated. Initially limited to Tasmania, this disease is currently causing major economic and fish welfare burden to Norwegian, Scottish and Australian salmon aquaculture (Shinn et al. 2015). The causative agent of this disease is the amoeba *Neoparamoeba perurans*, an opportunistic pathogen that can only be removed with expensive and laborious fresh water or hydrogen peroxide treatments (Novak and Archibald, 2018), and there are currently very limited opportunities for prevention.

A promising avenue to decrease the incidence of AGD in farmed Atlantic salmon is to increase genetic resistance of aquaculture stocks to *N*. *perurans*. There is significant genetic variation in resistance to AGD in commercial Atlantic salmon populations (Taylor et al. 2007, 2009; Kube et al. 2012; Robledo et al. 2018; Boison et al. 2019), therefore selective breeding has potential to improve gill health via a reduction in amoebic load and associated gill damage. The use of genetic markers through genomic selection can expedite genetic gain in aquaculture breeding programmes (e.g. Robledo et al. 2018; Yoshida et al. 2018; Palaiokostas et al. 2018a, 2018b), however, the need to genotype a large number of animals and to perform disease challenges in every generation involves a relatively high cost. The discovery of the mechanisms leading to resistance and the underlying causative genetic variants has the potential to reduce this cost via incorporation of functional SNPs into the genomic prediction models.

Discovering the genes and pathways that lead to successful immune responses to pathogens is a major goal in genetics and immunology research. Understanding disease resistance can aid selective breeding via incorporation of putative causative variants with greater weighting in genomic prediction models, which can improve selection accuracy and reduce the need for routine trait recording (MacLeod et al. 2016; Houston et al. 2017). Such information can also inform the development of improved disease challenge models, and more successful prevention or treatment strategies through an increased knowledge of host-pathogen interactions. Finally, with the potential role for targeted genome editing (e.g. using CRISPR/Cas9) in future aquaculture breeding programmes, understanding the functional mechanisms underlying disease resistance traits is key to identifying target genes and variants for editing. Previous studies into AGD-infected Atlantic salmon have suggested that the amoebae might elicit an immunosuppressive effect on the innate response of the host, with a concurrent up-regulation of the adaptive Th2-mediated response (Benedicenti et al. 2015; Marcos-López et al. 2017, 2018). Th2 cytokines were also found consistently up-regulated when comparing AGD infected and non-infected samples, and lesion and non-lesion areas (Marcos-López et al. 2018). The heritability of resistance to AGD has been shown to increase after successive cycles of disease challenge / treatment (Taylor et al. 2009), which could suggest that the ability to elicit a successful adaptive immune response is partly under genetic control. Finally, a higher expression of genes related to adaptive immunity has been previously reported in more AGD-resistant salmon compared to their more susceptible counterparts using a microarray approach to measure gene expression (Wynne et al. 2008).

In a previous study by our group, several QTL regions with a significant contribution to genetic AGD resistance were identified in Atlantic salmon derived from a commercial breeding programme (Robledo et al. 2018). In the current study, the gill and head kidney transcriptomes of AGD resistant and susceptible Atlantic salmon from the same population were sequenced and compared. The main goals of the study were a) to assess the differences in local and systemic immune responses between AGD resistant and susceptible Atlantic salmon, and b) to use gene expression data to identify positional and functional candidate genes underlying the previously detected resistance QTL.

## RESULTS

### Sampling and sequencing

Fish were classified into resistant or susceptible based on their mean gill score and their gill amoebic load. A previous study by our group has shown a high positive genetic correlation between these two traits (higher gill score associated with higher amoebic load), and both are considered indicator traits for resistance to AGD. RNA sequencing was performed on the gill and head kidney of 12 resistant and 12 susceptible fish. Resistant animals had a mean gill score of 2.92 ± 0.13, mean amoebic load (qPCR ct value) of 37.12 ± 3.63 and mean weight of 543 ± 116 g at the point of sampling; susceptible animals had a mean gill score of 4.12 ± 0.20, mean amoebic load of 25.99 ± 1.80 and mean weight of 409 ± 96 g. Sequencing of one of the gill samples rendered an extremely low number of reads and therefore was discarded. The remaining samples had an average of 24M filtered paired-end reads, which were pseudoaligned to genes in the Atlantic salmon genome. Exploratory analyses based on distance measures revealed two head kidney samples as outliers and they were removed (Additional file 1). Therefore, the final dataset comprised of 23 gill and 22 head kidney samples from 24 individuals. The two tissues showed clearly distinct patterns of gene expression, as would be expected. The difference in global gene expression pattern between resistant and susceptible samples in both tissues was much less pronounced, but still evident in the gill in particular (Figure 1). Similar results were described in a Norwegian commercial population (Boison et al. 2019).

**Figure 1.**
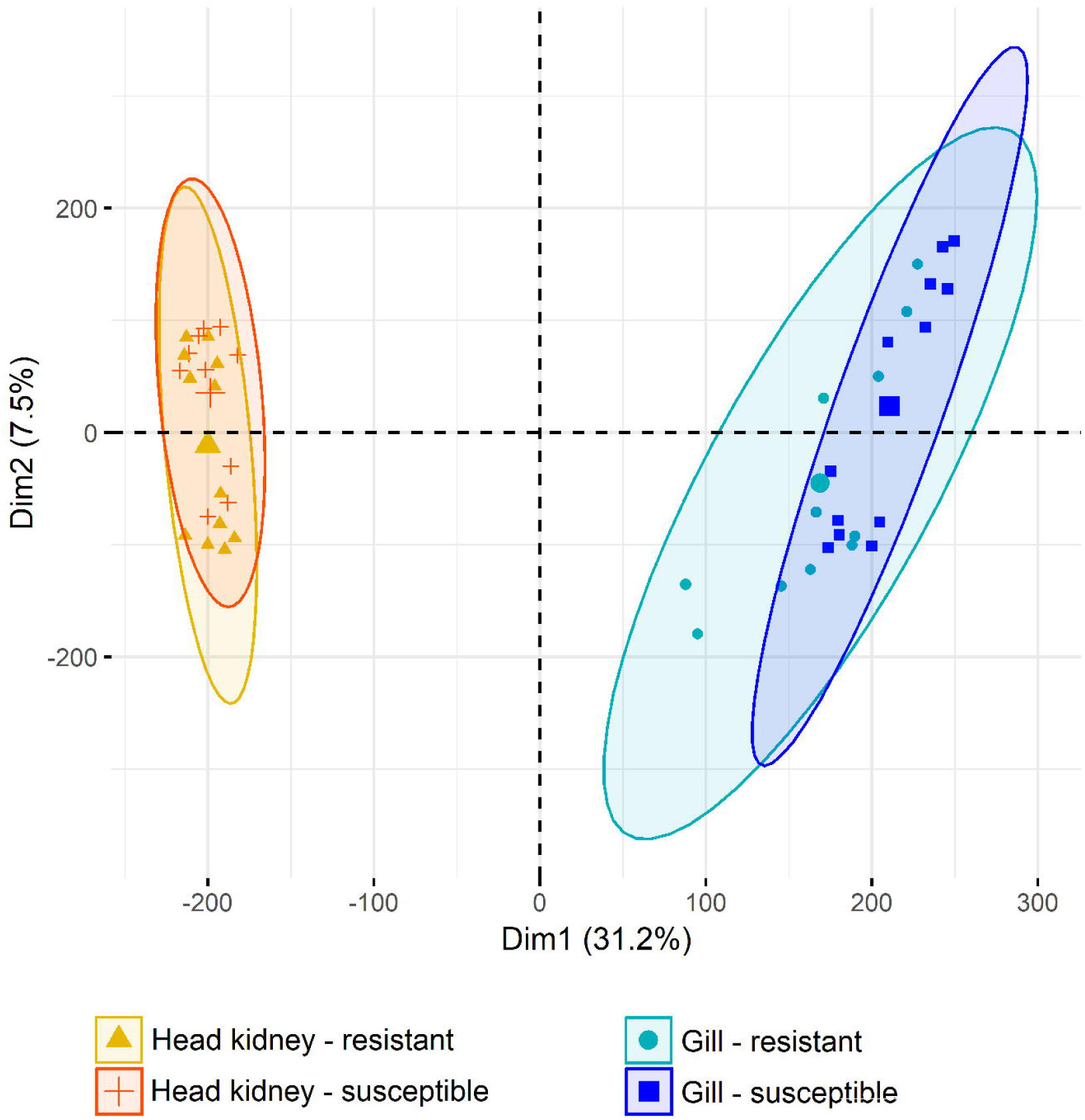
Principal component analysis. RNA-Seq samples clustered according to their gene expression. The larger symbols represent group means, and ellipses represent 95% confidence intervals for the groups.

### Differential expression

A total of 115 and 42 differentially expressed transcripts (following multiple-testing correction, FDR p-value < 0.05) were detected between resistant and susceptible samples in gill and head kidney tissues respectively (Figure 2, additional file 2). The clearest evidence for differential immune responses was found in gill, where several differentially expressed immune-related transcripts were detected. Most differentially expressed transcripts in head kidney were not obviously related to AGD or disease resistance. To gain an overall view of the results we performed a gene Ontology (GO) enrichment test in both gill and head kidney for sets of differentially expressed transcripts according to three different significance criteria (p-value < 0.01, 0.05 and 0.1) (Figure 3). In the gill, various relevant GO terms were observed, such as “Response to stress”, “Cytoskeleton” and “Circulatory system process”. We observed a larger number of enriched GO terms in head kidney. While most of them are seemingly unconnected to AGD resistance, terms such as “Response to stress” or “Protein modification process” were observed. For instance, of 22 genes showing p-values < 0.01, 15 of them were assigned to “Response to stress”. Similar analyses for KEGG pathways did not reveal any significant enrichment.

**Figure 2.**
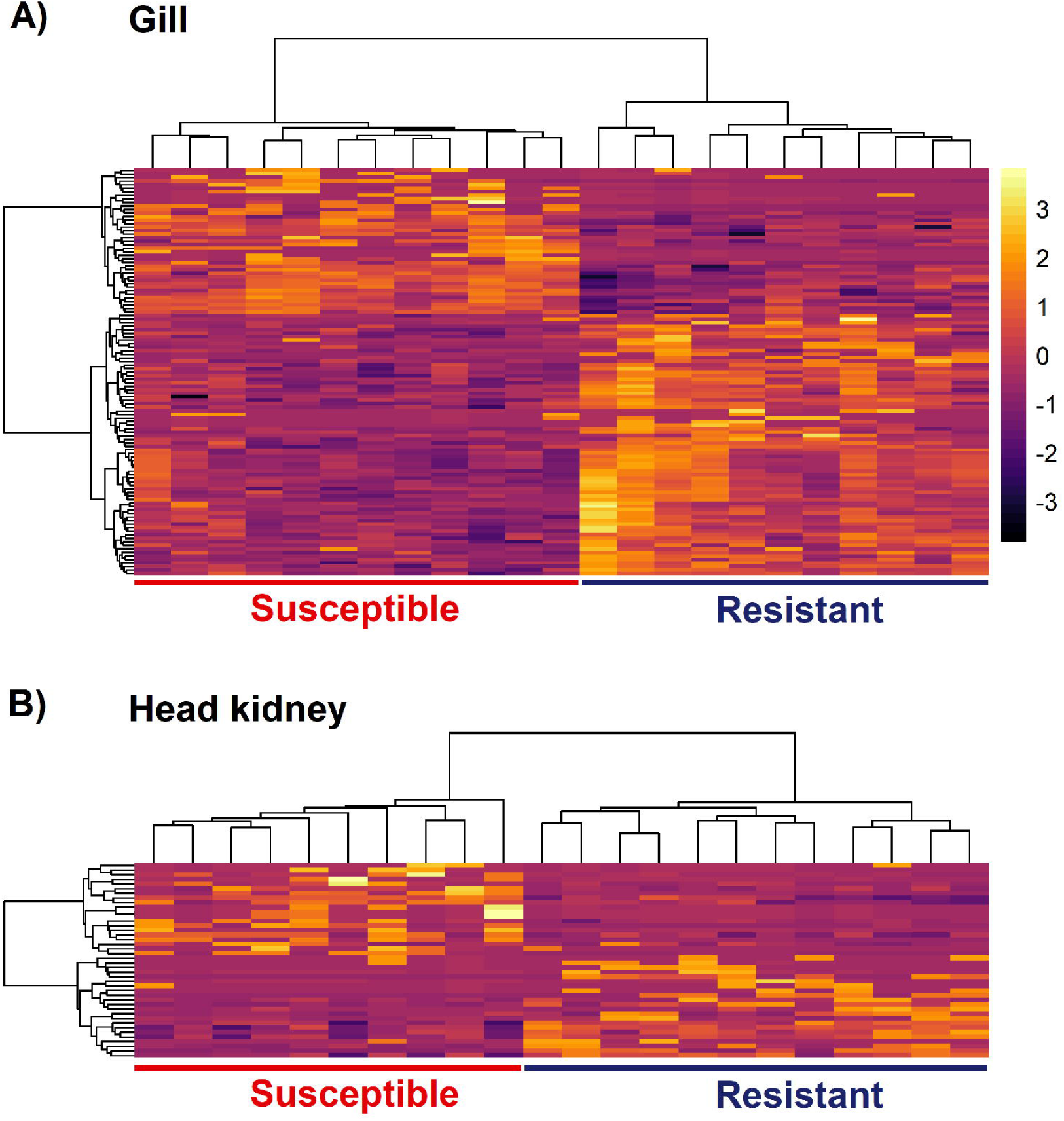
Heatmap of differentially expressed genes between resistant and susceptible samples. Heatmaps of all differentially expressed genes in gill (A) and head kidney (B). Samples and genes were clustered according to gene expression (mean centered and scaled normalized counts).

**Figure 3.**
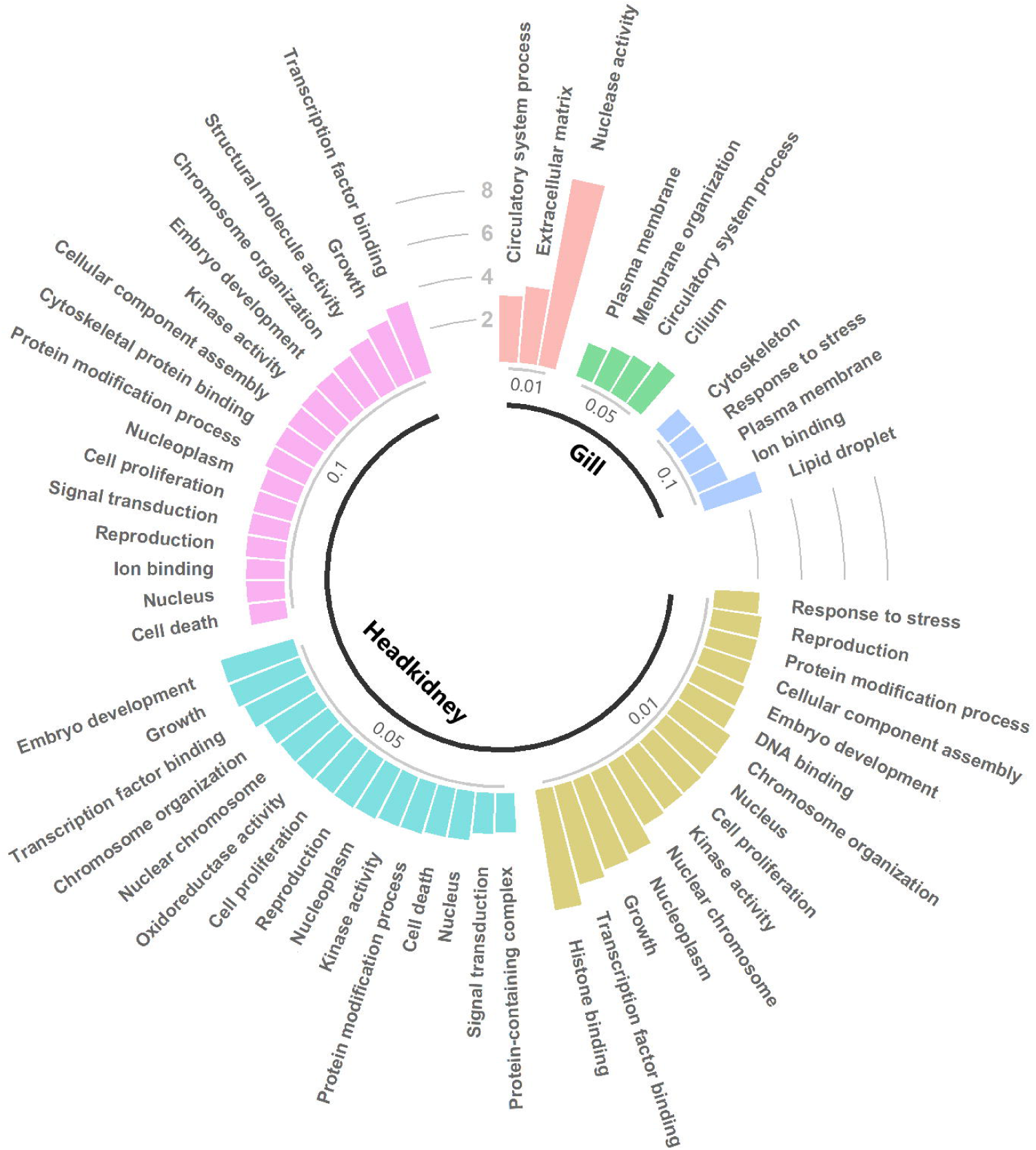
Gene Ontology enrichment for differentially expressed genes. GO enrichment is shown for all differentially expressed genes in gill and head kidney according to three different significant criteria (FDR p-value < 0.1, 0.05 and 0.01). The height of the bars represents fold enrichment (percentage of genes assigned to the GO term in the set of differentially expressed genes compared to the percentage assigned to that GO term in the transcriptome of that tissue).

Detailed inspection of the differentially expressed transcripts in the gill revealed that they can be grouped into three broad categories concordant with GO enrichment results: 1) immune response (“Response to stress”), 2) cell adhesion or cell shape (“Cytoskeleton”), and 3) red blood cells and coagulation (“Circulatory system process”).

Amongst the immune-related transcripts showing differential expression in the gill was interleukin-17 receptor E (*IL17RE*), which was highly expressed in resistant animals (logFC = 1.1). In mice *IL17RE* is the receptor for *IL-17C*, which has an essential role in host mucosal defense against infection and is critical for a successful immune response against bacterial infection (Song et al. 2011). The *IL-17C* – *IL17RE* pair also stimulates T-helper cell 17 responses, which has a proinflammatory effect (Chang et al. 2011). Benedicenti et al. (2015) reported that *IL-17C* expression showed a negative correlation with amoebic load in Atlantic salmon, and that the Th17 pathway in general was found to be significantly down-regulated in response to AGD. This could be a mechanism of immune evasion elicited by the parasite, which might be more effective in susceptible fish than resistant. Another highly expressed transcript is involved in T-cell function, T-cell specific surface glycoprotein CD28 (*CD28*; logFC = 1.60). *CD28* promotes T-cell survival and proliferation, and enhances the production of multiple cytokines including IL4 (Blotta et al. 1996) *IL4* has been found to be up-regulated in response to AGD (Marcos-López et al. 2018), and this gene induces differentiation of naïve helper T cells to Th2 cells. The Th2 pathway was found to be up-regulated in late stages of AGD (Benedicenti et al. 2015). This pathway is linked to humoral immune responses against extracellular parasites and to tissue repair (Allen and Sutherland, 2014), and therefore is an expected response to AGD. A higher prevalence of this type of response in resistant animals would also be consistent with the observed increase of the heritability of resistance after successive cycles of disease challenge / treatment (Taylor et al. 2009), reflecting genetic variability in the effectiveness of the adaptive response, and / or variation in immune memory.

Several genes connected to red blood cells were found to be differentially expressed, including five different haemoglobin subunit transcripts, which were highly expressed and clearly up-regulated in resistant samples in the gill (logFC ∼ 2). Reduced hematocrit has been described in AGD infected Atlantic salmon, linked mainly to gill damage (Hvas et al. 2017). The plasma protease C1 inhibitor gene (*SERPING1*) is also up-regulated (logFC = 1.2). This gene inhibits the complement system and also has anti-inflammatory functions (Davis et al. 2008). Complement proteins have been found in gill mucus of AGD infected Atlantic salmon (Valdenegro-Vega et al. 2014). The lower expression of *SERPING1* in susceptible samples might simply be a reflection of the higher extent of gill damage in these animals, requiring activation of the complement system and increase of local inflammatory responses.

There are also a few differentially expressed transcripts connected to cell adhesion and cell shape, including a cadherin gene (cadherin-related family member 5; logFC = 4.5) and an actin related gene (actin filament associated protein 1-like 1; logFC = 1.3). The Cdc42 effector protein 2 (*CDC4EP2*; logFC = 0.6) was also up-regulated in resistant fish, and has been associated with roles in actin filament assembly and control of cell shape (Hirsch et al. 2001). A previous study identified an enrichment of cell-adhesion genes in severely affected animals compared to others with healthier gills infected by AGD (Boison et al. 2019). These changes are consistent with the epithelial hyperplasia and other structural changes caused by the parasite in the gill of infected animals (Nowak 2012).

Head kidney differential expression results are much harder to interpret, and most of the DE transcripts are seemingly not connected to biological processes that have previously been related to AGD. Tumor necrosis factor alpha-induced protein 8-like protein 1 (*TNFAIP8L1*; logFC = -0.9) was found to be more highly expressed in susceptible samples. This gene inhibits apoptosis by suppressing the activity of caspase-8 (You et al. 2001). The down-regulation of pro-apoptotic genes has been connected to AGD severity (Marcos-López et al. 2018). The lack of a clear picture in head kidney might reflect the relative importance of the local and systemic immune responses in response to AGD. Two transcripts showed differential expression in both tissues: putative ferric-chelate reductase 1 (*FRRS1*) and CG057 protein.

The regulation of transcripts upon infection is a strong indication of the involvement of the gene product in the immune and physiological response of the host to the pathogen, but a comparison between resistant and susceptible animals can offer insight into the mechanisms determining the success of the immune response against the pathogen. The main caveat of this approach is that it is difficult to distinguish cause and consequence, i.e. is the gene differentially expressed because it confers resistance or due to differential disease progression? Additional evidence, such as the co-localization of differentially expressed genes with QTL or the identification of cis regulatory variants in the QTL regions can further contribute to our understanding of disease resistance, and help us discover underlying candidate genes.

### Integration with previous QTL

The overlap between previously identified QTL regions in this population (Robledo et al. 2018) and the differentially expressed genes was explored (Figure 4). A differentially expressed gene, interferon-induced very large GTPase 1 (*GVINP1*), was found in one of the QTL regions of chromosome 18, which explained ∼20% of the genetic variance in resistance to AGD (second largest QTL). Very little is known about the function of this gene, but it has been shown to respond to both type I and type II interferon response in mammals (Klamp et al. 2003). The genes showing FDR corrected p-values < 0.1 (a total of 268 genes) were also investigated, and four additional genes were found in these QTL regions. *MAP4K4*, located in a putative QTL region of chromosome 17, surpassed this threshold, and is involved in systemic inflammation in mammals (Aouadi et al. 2009), and *TRIM39* in the second QTL region in chromosome 18, a positive regulator of apoptosis (Rosenthal 2012).

**Figure 4.**
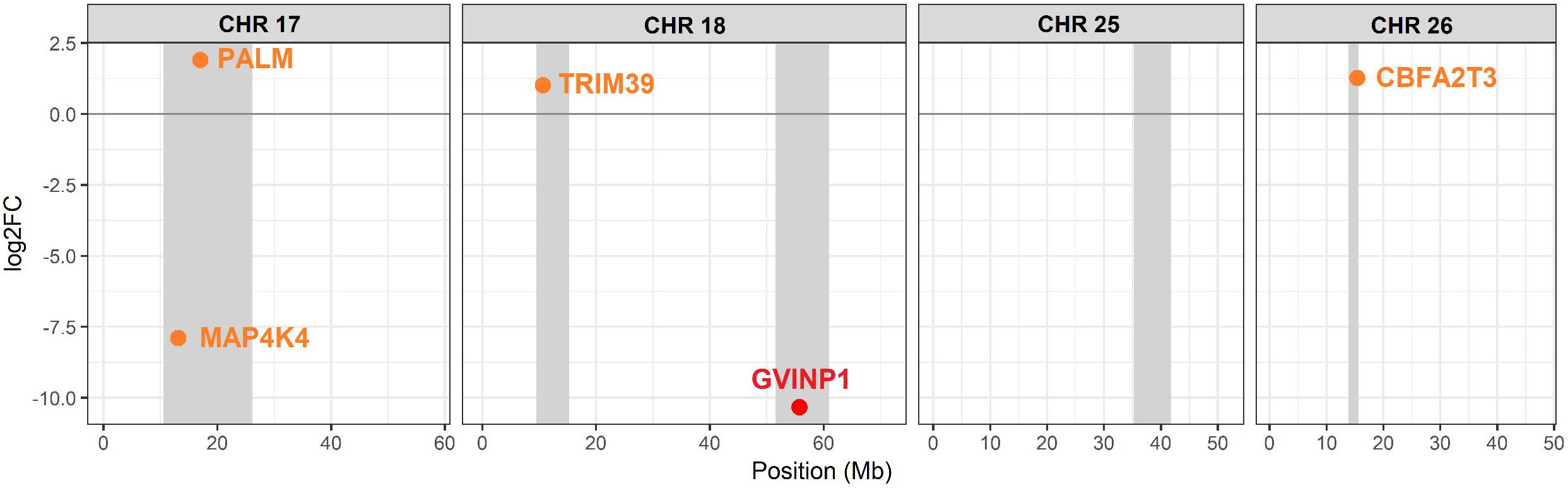
Differentially expressed genes located in resistance QTL. The location of the QTL regions in the chromosomes are shown in grey. Genes with significance values < 0.05 are in red, those with significance values < 0.1 are in orange. Positive fold changes correspond to higher expression in resistant samples.

**Figure 5.**
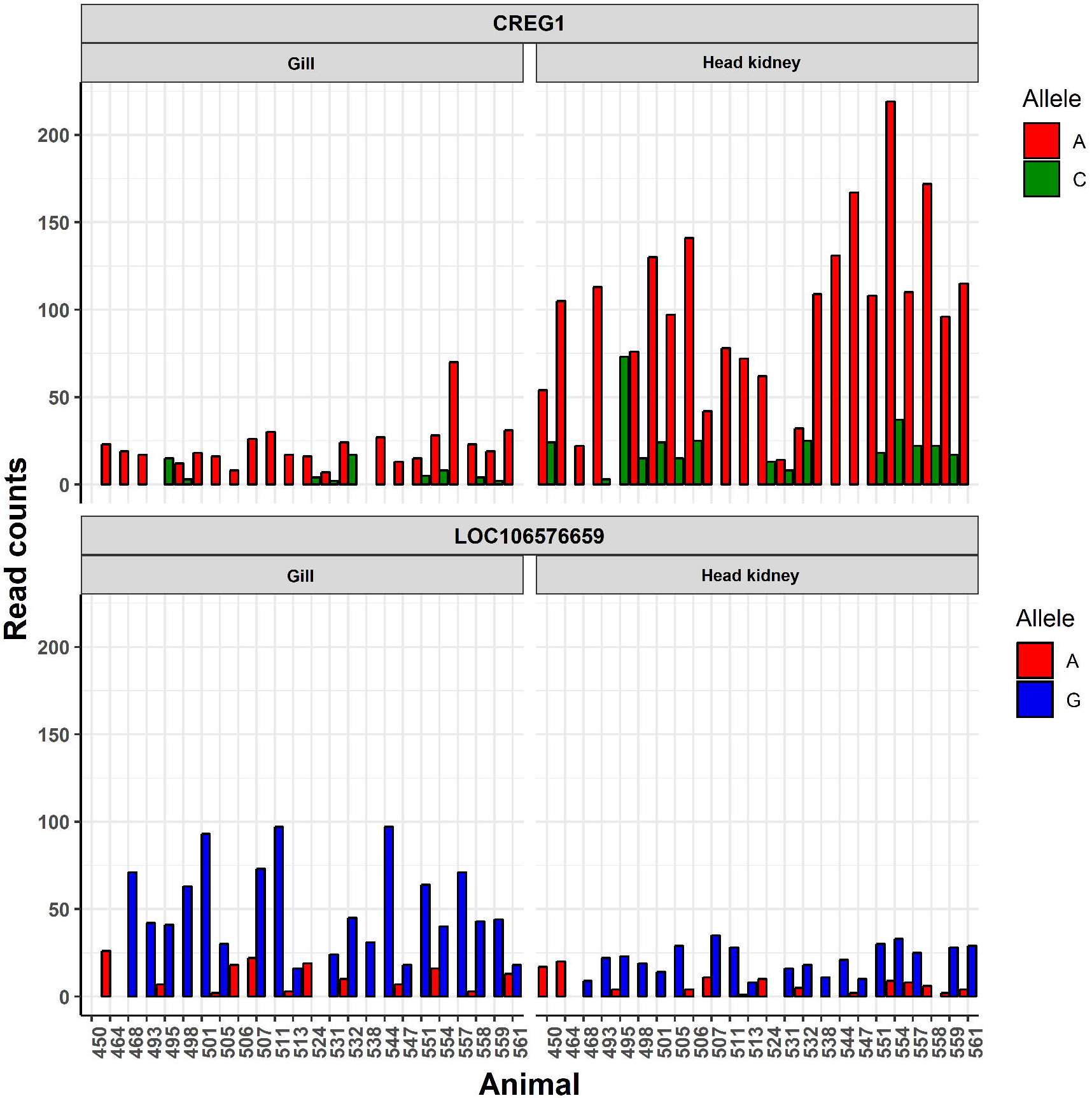
Allele specific expression CREG1. Barplot showing the read counts for each allele for those SNPs in the QTL regions showing allele specific expression. The two SNPs are located in *CREG1* (Chromosome 17 – 24,545,527 bp) and the uncharacterized gene LOC106576659 (Chromosome 18 – 57,163,493 bp).

### Allele specific expression

To explore potential cis-acting variation underlying the resistance QTL, an allele specific expression (ASE) test was performed for the SNPs in transcripts within the QTL regions, finding a significant ASE event in a gene in chromosome 17; cellular repressor of E1A-stimulated genes 1 (*CREG1*). In humans this protein is connected to the regulation of cellular proliferation and differentiation (Di Bacco and Gill, 2003), and antagonizes the proliferative effects of adenovirus E1A protein (Veal et al. 1998). This gene showed a log fold change of 0.75 between resistant and susceptible samples in gill (FDR p-value = 0.25). A second significant ASE event was found in an uncharacterised gene in chromosome 18 (LOC106576659), however this gene showed no differences in fold change between resistant and susceptible samples.

The polygenic nature of resistance to AGD means that different resistance mechanisms might be operating in each different family. The connection between genotypes and expression, through expression QTL (eQTL) or ASE, can provide strong evidence for functional candidate genes underlying QTL. While eQTL studies require a relatively large number of animals, the advantage of ASE is that the statistical test in performed separately in each heterozygous individual. It is well known that most causative variants are part of regulatory elements and affect gene expression (Keane et al. 2011; Albert and Kruglyak 2015), therefore the detection of ASE in a QTL can provide strong evidence linking the function of a gene to the QTL and the phenotype of interest.

## DISCUSSION

The potential benefits of the identification of causative variation impacting on complex traits are substantial, ranging from fundamental knowledge of the biology underlying the traits of interest to their application for enhancing these traits in farmed populations. However, even with the addition of various layers of information such as RNA sequencing, determining the causative gene underlying a QTL is not straightforward, especially because the QTL regions tend to be large and contain a large number of genes, as previously described for sea lice resistance QTL (Robledo et al. 2019).

Eventually, functional assays are necessary to provide actual evidence of its causality. The advent of CRISP-CAS9 has made this much more feasible in non-model species. Likewise, this technology now provides the opportunity of using this information to introduce or fix favorable alleles in farmed populations. The genetic architecture of quantitative traits usually varies across populations, and indeed AGD resistance QTL seem to vary across different Atlantic salmon commercial populations (Robledo et al. 2018, Boison et al. 2019). While the use of genome editing in farmed animals requires societal and regulatory changes, the transference of causative variants across populations can lead to a rapid increase in disease resistance (Jenko et al. 2015), with long-lasting effects on animal welfare and food security. Nevertheless, the discovery of causative variants and genes can be used to increase the weight of causative variants in genomic selection, increasing its accuracy and therefore speeding up genetic gain in each generation (MacLeod et al. 2016). More widely, basic knowledge about the pathways leading to resistance to disease can inform drug development or preventive measures such as functional feeds. To summarise, finding the underlying cause(s) of resistance to disease can provide large benefits for aquaculture and society in the form of healthier animals, increased food security and sustainable economic gain, directly through their implementation in breeding schemes in the present and through genome editing in the future.

## CONCLUSIONS

The transcriptomic differences between AGD resistance and AGD susceptible Atlantic salmon are limited, which might not be surprising considering the polygenic nature of the trait. The differences were more evident in the gill than in head kidney, potentially highlighting the importance of the local immune response. Genes involved in immune response (Th2 and Th17 pathways), red blood cells and cell adhesion could be part of the mechanisms leading to AGD resistance, however, it is difficult to discriminate cause and consequence. The integration of previously discovered QTL and expression data pointed to potential candidate genes of interest, such as *GVINP1, MAP4K4* or *TRIM39*. An additional candidate gene, *CREG1*, showed allele specific expression in one of the QTL regions. Follow-up studies to investigate the functional role of these genes in the response to AGD could help us understand the molecular mechanism of resistance to this parasite, and contribute to improving gill health in farmed populations through incorporation of functional data to improve genomic prediction, or potentially via genome editing in the longer term.

## METHODS

### Experimental design

The AGD challenge experiment was performed using 797 Atlantic post-smolt salmon from 132 nuclear families (∼18 months, mean weight after challenge ∼464 g) originating from a commercial breeding programme (Landcatch, UK). The challenge experiment was performed as described in Robledo et al. (2018). In brief, seeder fish with a uniform level of AGD infection were produced by cohabitation with fish infected from an *in vivo* culture. The challenge was then performed by cohabitation of infected seeder fish at a ratio of 15% seeder to naïve fish, allowing three separate cycles of infection with a treatment and recovery period after the first two (Taylor *et al*. 2009) using a 4 m^3^ seawater tank in the experimental facilities of University of Stirling’s Marine Environmental Research Laboratory, Machrihanish (Scotland, UK). The fish were kept under a 16-h light and 8-h dark photoperiod, starting at 05:00; the fish were fed Biomar organic salmon feed, automatic every 20 minutes to approximately 1% biomass; water supply was ambient flow-through filtered to approximately 90 microns, for the duration of the trial, water temperature was between 13 & 14 °C and salinity was 33-35ppt. For the first two challenges, fresh water treatment was performed 21 days after challenge, followed by a week of recovery. The fish were checked visually four times daily during this period. The disease was allowed to progress until the terminal sampling point in the third challenge, when fish were terminated by an overdose of anaesthetic followed by destruction of the brain. Fish were sampled and phenotypes were recorded during three consecutive days. A subjective gill lesion score of the order of severity ranging from 0 to 5 was recorded for both gills (Taylor *et al*. 2016). These gill lesion scores were recorded by a single operator, who referred to pictures to guide classification. Some fish were scored by additional operators, and the scores never differed by > 0.5. Further, one of the gills was stored in ethanol for qPCR analysis of amoebic load using *N*. *perurans* specific primers. Amoebic load has previously been used as a suitable indicator trait for resistance to AGD in salmon (Taylor *et al*. 2009). All fish were phenotyped for mean gill score (mean of the left gill and right gill scores) and amoebic load (qPCR values using *N*. *perurans* specific primers, amplified from one of the gills). 24 fish from 24 different families were selected for RNA sequencing (Additional file 3) based on high or low levels of resistance according to the measured traits (mean gill score and amoebic load as measured by qPCR). Gill and head kidney samples were obtained from each animal and stored in RNAlater at 4 °C for 24 h, and then at -20°C until RNA extraction.

### RNA extraction and sequencing

For all the 48 samples a standard TRI Reagent RNA extraction protocol was followed. Briefly, approximately 50 mg of tissue was homogenized in 1 ml of TRI Reagent (Sigma, St. Louis, MO) by shaking using 1.4 mm silica beads, then 100 µl of 1-bromo-3-chloropropane (BCP) was added for phase separation. This was followed by precipitation with 500 µl of isopropanol and subsequent washes with 65-75 % ethanol. The RNA was then resuspended in RNAse-free water and treated with Turbo DNAse (Ambion). Samples were cleaned up using Qiagen RNeasy Mini kit columns and their integrity was checked on Agilent 2200 Bioanalyzer (Agilent Technologies, USA). Thereafter, the Illumina Truseq mRNA stranded RNA-Seq Library Prep Kit protocol was followed directly. Libraries were checked for quality and quantified using the Bioanalyzer 2100 (Agilent), before being sequenced on three lanes of the Illumina Hiseq 4000 instrument using 75 base paired-end sequencing at Edinburgh Genomics, UK. Raw reads have been deposited in NCBI’s Sequence Read Archive (SRA) under BioProject accession number PRJNA552604.

### Read mapping

The quality of the sequencing output was assessed using FastQC v.0.11.5 (http://www.bioinformatics.babraham.ac.uk/projects/fastqc/). Quality filtering and removal of residual adaptor sequences was conducted on read pairs using Trimmomatic v.0.38 (Bolger et al. 2004). Specifically, Illumina specific adaptors were clipped from the reads, leading and trailing bases with a Phred score less than 20 were removed and the read trimmed if the sliding window average Phred score over four bases was less than 20. Only reads where both pairs were longer than 36 bp post-filtering were retained. Filtered reads were mapped to the most recent Atlantic salmon genome assembly (ICSASG_v2; Genbank accession GCF_000233375.1; Lien et al. 2016) using STAR v.2.6.1a (Dobin et al. 2013), the maximum number of mismatches for each read pair was set to 10 % of trimmed read length, and minimum and maximum intron lengths were set to 20 bases and 1 Mb respectively. Transcript abundance was quantified using kallisto v0.44.0 (Bray et al. 2016) and the latest Atlantic salmon genome annotation (NCBI *Salmo salar* Annotation Release 100).

### Differential Expression

Differential expression (DE) analyses were performed using R v.3.5.2 (R Core Team 2014). Gene count data were used to estimate differential gene expression using the Bioconductor package DESeq2 v.3.4 (Love et al. 2014). Briefly, size factors were calculated for each sample using the ‘median of ratios’ method and count data was normalized to account for differences in library depth. Next, gene-wise dispersion estimates were fitted to the mean intensity using a parametric model and reduced towards the expected dispersion values. Finally a negative binomial model was fitted for each gene and the significance of the coefficients was assessed using the Wald test. The Benjamini-Hochberg false discovery rate (FDR) multiple test correction was applied, and transcripts with FDR < 0.05 and absolute log_2_ fold change values (FC) > 0.5 were considered differentially expressed genes. Hierarchical clustering and principal component analyses were performed to visually identify outlier samples, which were then removed from the analyses. PCA plots were created using the R package factoextra (http://www.sthda.com/english/rpkgs/factoextra/).

Gene Ontology (GO) enrichment analyses were performed in R v.3.5.2 (R Core Team 2014) using Bioconductor packages GOstats v.2.48.0 (Falcon and Gentleman, 2007) and GSEABase v.1.44.0 (Morgan et al. 2018). GO term annotation for the Atlantic salmon transcriptome was obtained using the R package Ssa.RefSeq.db v1.3 (https://gitlab.com/cigene/R/Ssa.RefSeq.db). The over-representation of GO terms in differentially expressed gene lists compared to the corresponding transcriptomes (gill or head kidney) was assessed with a hypergeometric test. GO terms with ≥ 5 DE genes assigned and showing a p-value < 0.05 were considered enriched. Kyoto Encyclopedia of Genes and Genomes (KEGG) enrichment analyses were performed using KOBAS v3.0.3 (Wu et al. 2006). Briefly, salmon genes were annotated against KEGG protein database (Kanehisa and Goto 2000) to determine KEGG Orthology (KO). KEGG enrichment for differentially expressed gene lists was tested by comparison to the whole set of expressed genes in the corresponding tissue using Fisher’s Exact Test. KEGG pathways with ≥ 5 DE genes assigned and showing a Benjamini-Hochberg FDR corrected p-value < 0.05 were considered enriched for differential expression. The reference tissue transcriptome for both GO and KEGG enrichment comprised only those genes with mean normalized counts value > 5.

### Allele specific expression

Gene expression estimates and genotypes obtained from the RNA sequencing were used to investigate allele specific expression. The samtools v1.6 software (Li et al. 2009) was used to identify SNPs, and call genotypes for those SNPs in individual samples. PCR duplicates, reads with mapping quality < 20 and bases with phred quality scores < 20 were excluded. SNPs within 5 bp of an indel, with quality < 20, MAF < 0.05 or less than 4 reads supporting the alternative allele were discarded. The putative effect of the SNPs was assessed using the official salmon genome annotation (NCBI *Salmo salar* Annotation Release 100) and the SnpEff v.4.2 software (Cingolani et al. 2012). Allelic specific expression was assessed using the R package AllelicImbalance (Gadin et al. 2015). For every SNP in the regions of interest, read counts were obtained for each allele in heterozygous animals, those with less than 10 reads were filtered, and a binomial test was performed to assess the significance of the allelic differences. Only those genomic positions called as heterozygotes in a minimum of 4 and a maximum of 36 (75%) samples (75%) were considered. An allele specific expression event was considered significant if the mean p-value of all heterozygotes was < 0.05. All significant events were manually inspected.

## Supporting information

Addtional file 2

## DECLARATIONS

### Ethics approval and consent to participate

All animals were reared in accordance with relevant national and EU legislation concerning health and welfare. The challenge experiment was performed by the Marine Environmental Research Laboratory (Machrihanish, UK) under approval of the ethics review committee of the University of Stirling (Stirling, UK) and according to Home Office license requirements. Landcatch are accredited participants in the RSPCA Freedom Foods standard, the Scottish Salmon Producers Organization Code of Good Practice, and the EU Code-EFABAR Code of Good Practice for Farm Animal Breeding and Reproduction Organizations.

## Consent for publication

Not applicable

## Availability of data and materials

The RNA sequencing dataset generated during the current study is available in NCBI’s Sequence Read Archive (SRA) under BioProject accession number PRJNA552604. The phenotypes of all sequenced samples are included in this published article (Additional file 3).

## Author’s contributions

RH and DR were responsible for the concept and design of this work and drafted the manuscript. AH was responsible for the disease challenge. AH and JB managed the collection of the samples. AG performed the molecular biology experiments. DR performed bioinformatic and statistical analyses. All authors read and approved the final manuscript.

## Acknowledgements

Not applicable

## ADDITIONAL FILES

### Additional file 1

Portable Network Graphics (.png)

Principal component analysis of all RNA sequenced samples

RNA-Seq samples clustered according to their gene expression. Outliers were discarded for further analyses.

### Additional file 2

Excel file (.xlsx)

Differentially expressed genes between resistant and susceptible samples

Lists of differentially expressed genes between resistant and susceptible samples in gill and head kidney. Gene ID, position in the Atlantic salmon genome (Chromosome, start and end in base pairs), average expression of the gene, log 2 fold change between resistant and susceptible animals (positive fold changes correspond to higher expression in resistant samples), standard deviation of the fold change, FDR adjusted p-value, gene annotation and gene symbol are shown.

### Additional file 3

Excel file (.xlsx)

Phenotypes of all samples used for RNA sequencing

All collected phenotypes for the samples used in this study. ID of the sample, tissue, whether it is resistant or susceptible, finclip ID linking to the genotypes (available in Robledo et al. 2018), gill scores for both gills (and mean), weight and length at the end of the challenge, and amoebic load measured by qPCR are shown.

