## Supplementary figures and images for "Characterising the mechanisms underlying genetic resistance to amoebic gill disease in Atlantic salmon using RNA sequencing"

### Additional file 1

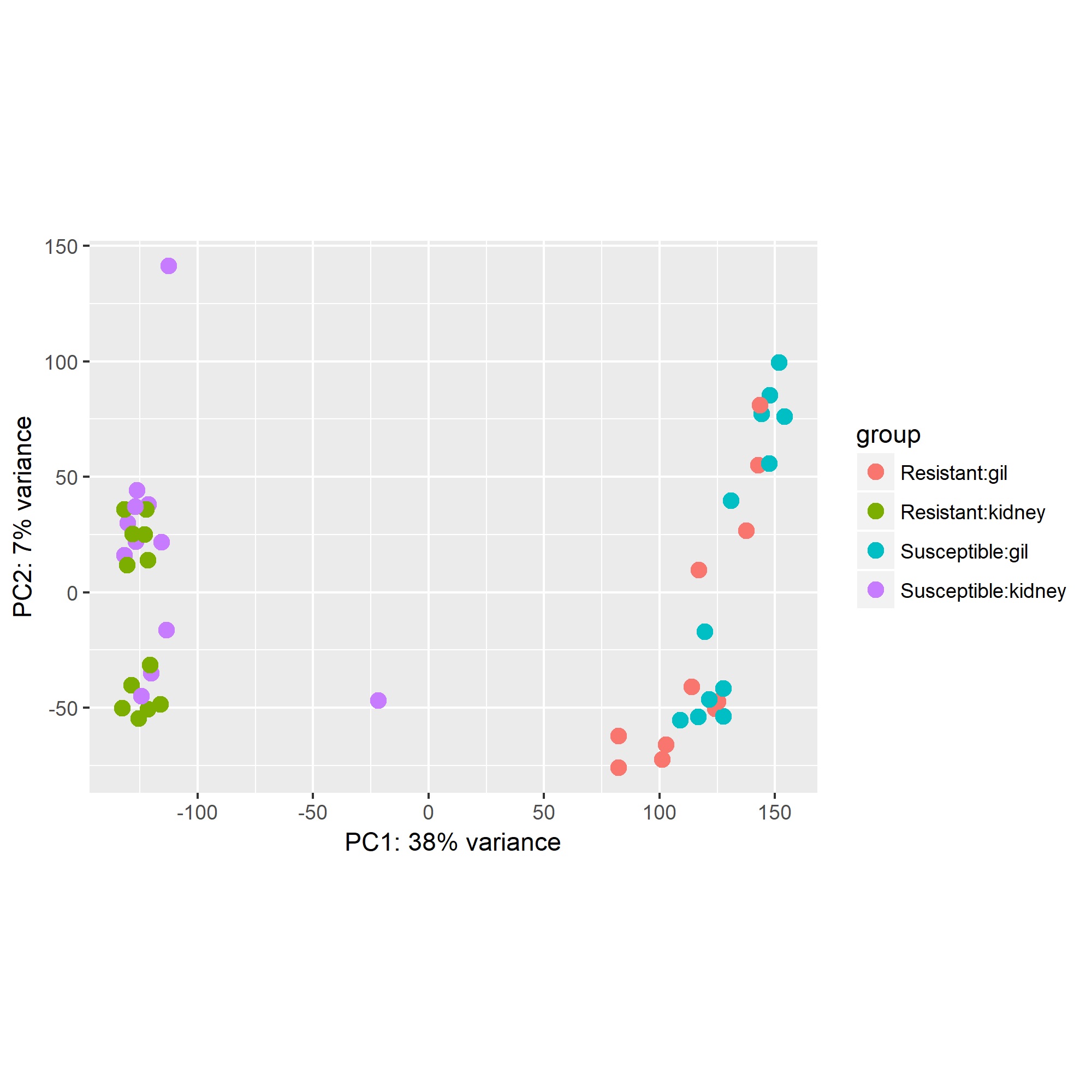
